# STAR Chimeric Post For Rapid Detection of Circular RNA and Fusion Transcripts

**DOI:** 10.1101/139808

**Authors:** Nicholas K. Akers, Eric E. Schadt, Bojan Losic

**Affiliations:** Department of Genetics and Genomic Sciences, The Icahn Institute for Genomics and Multiscale Biology Icahn School of Medicine at Mount Sinai, One Gustave L. Levy lace, New York, NY 10029, USA

## Abstract

**Motivation:** The biological relevance of chimeric RNA alignments is now well established. Chimera arising as chromosomal fusions are often drivers of cancer, and recently discovered circular RNA are only now being characterized. While software already exists for fusion discovery and quantitation, high false positive rates and high run-times hamper scalable fusion discovery on large datasets. Furthermore, very little software is available for circular RNA detection and quantification.

**Results:** Here we present STAR Chimeric Post (STARChip), a novel software
package that processes chimeric alignments from the STAR aligner and produces annotated circular RNA and high precision fusions in a rapid, efficient, and scalable manner that is appropriate for high dimensional medical omics datasets.

**Availability and Implementation:** STARChip is available at https://github.com/LosicLab/STARChip

**Supplementary Information:** Supplementary figures and tables are available online.

## 1. Introduction

The key agnostic hallmark of the RNA-Seq assay compared to microarray is the potential to observe previously unknown RNA fragments. This revolutionary power, in principle, allows for a complete de-novo sampling of the transcriptome. At present, however, this ideal is rarely attained in practice. Difficulties of computation, interpretation, and validation typically impede one from attempting to leverage RNA-Seq beyond straightforward linear gene expression analysis. Nevertheless, confronting the reality of the complicated, dynamically spliced eukaryotic transcriptome in large high dimensional omics datasets naturally raises important and increasingly tractable questions about non-mRNA fragments, including circular RNA (circRNA) and RNA from chromosomal rearrangements.

In fact, there is a rapidly growing field of research indicating that circular isoforms of RNA are common, tissue specific(Salzman *et al.*, 2013), expressed across eukaryotes(Wang *et al.*, 2014) and may be associated with disease(Bachmayr-Heyda *et al.*, 2015). The molecular function of circRNA is unknown, with evidence indicating circRNA can regulate microRNA(Hansen *et al.*, 2013), though this is not likely the function of most circRNA(Guo *et al.*, 2014). CircRNA lack polyA tails, and can be detected in RNA that has been prepared using a RiboZero protocol (Figure 1). Perhaps surprisingly, for at least a fraction of genes accurate quantification of protein coding transcripts can be confounded by circRNA abundance(Salzman *et al.*, 2013).

**Figure 1.**
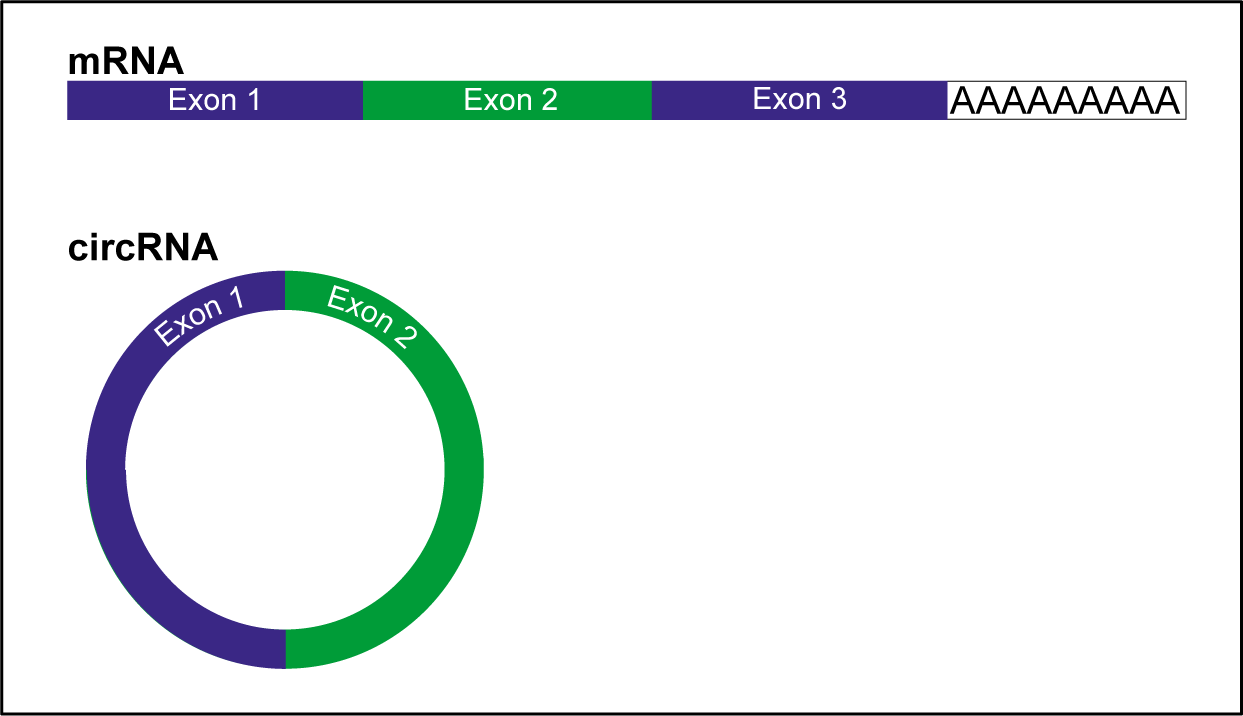
Description of typical circRNA compared to mRNA. mRNA (top) are typically composed of several exons and a poly-adenine tail. CircRNA (bottom) are commonly composed of exons, however they lack poly-adenine tails.

Non-linear RNA alignments can also be used to detect chromosomal rearrangements, a common causal factor in cancer. Chromosomal fusions are the aberrant connection of part of one chromosome with another. Initially described in chronic myelogeneous leukemia, recurrent chromosomal fusions have been described in 20 types of cancer(Mertens *et al.*, 2015). Uncovering these fusions contributes to knowledge of the pathogenesis of disease as well as serving as clinical biomarkers. Because chromosomal fusions may occur in non-transcribed regions of the genome, RNA-Seq is limited in its ability to observe all such events. On the other hand, whole genome sequencing is relatively expensive and does not generally provide any information about gene expression.

The landscape of software tools for circRNA and RNA fusion detection is actively evolving. For the rapidly developing field of circRNA, the existing options generate results that are inconsistent with one another(Hansen *et al.*, 2016), and only one software package (CIRI (Gao *et al.*, 2015)) leverages the power of multiple samples to improve circRNA prediction. Fusion detection is a more developed field with several mature software options(Jia *et al.*, 2013; STAR-Fusion/STAR-Fusion; Nicorici *et al.*, 2014; Kim and Salzberg, 2011), however we will show that these tools suffer from a high false-positive rate that prohibits validation using limited patient DNA. Additionally, the majority of existing fusion detection software packages perform alignments as a part of their sequence alignment pipeline, which prevents the use of the same alignments for both chimera detection and linear gene expression quantification. This can represent a significant increase in the computational burden of any bioinformatics pipeline.

Reasoning that a simplified filtration of high quality chimeric alignments will improve circRNA and fusion detection, we created a single software package based on the STAR aligner(Dobin *et al.*, 2013), STAR Chimeric Post (STARChip).

## 2. Methods

STARChip is written in Perl, Bash, and R. It is implemented in two distinct modes; detection of circRNA or fusion transcripts (Figure 2). STARChip is able to process raw sequence (FASTQ format) or to directly use the output of the STAR aligner. Although the ideal parameters will vary with desired sensitivity and the length of the reads, in many cases the STAR flags, “–chimSegmentMin 15 –chimJunctionOverhangMin 15” can be used to generate high quality chimeric output.

**Figure 2.**
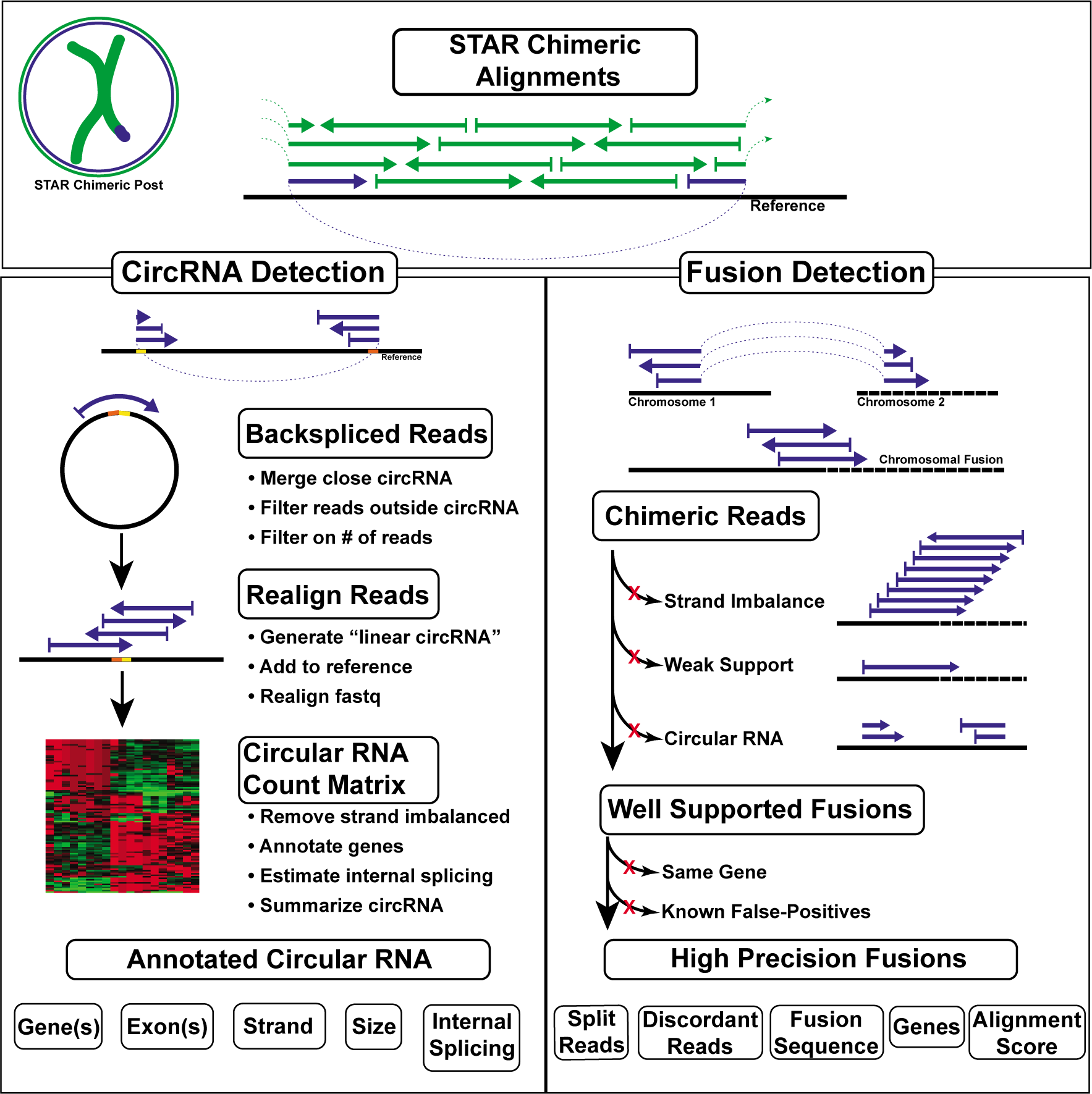
STARChip Flow Diagram. CircRNA (left side) are processed by filtering STAR Chimeric output files to discover backspliced reads, which align on the same chromosome and strand, in a non-linear manner. These reads are proximity-merged with other circRNA, filtered if part of the read aligns outside the circRNA, and summarized for each sample. Filtering on the number of reads and sample frequency provides a list of circRNA for follow-up. A count matrix of circRNA in each sample is generated, and summary visualizations are generated. Optionally, genes are annotated for each end of the circRNA, and linear splicing internal and external to the circRNA is summarized. Several tables are produced, which conveniently summarize the circRNA for follow-up. Fusion transcripts (right side) are identified by first collapsing all chimeric reads on location. Those putative fusion sites with strand imbalanced support (ie aligned reads are selectively aligned to a single strand), very weak support, or that appear to be circRNA are excluded. Fusions are then annotated for genes, known CNVs, and repeat regions. Fusions within the same gene or CNV are excluded, as are fusions that fall into known false-positive gene pairs. Finally, well-annotated, high precision fusions are written in tab-delimited format.

It is recommended to use a STAR index containing all reference chromosomes as well as unplaced contigs—reads transcribed from unplaced contigs may map chimerically without a proper reference. For annotation, STARChip relies on reference gene annotation (GTF) and sequence (FASTA) files, which must be modified for use with STARChip using a built in script. Software dependencies are R(R Core Team, 2015), BEDTools(Quinlan, 2014), SAMtools(Li *et al.*, 2009), and MAFFT(Katoh and Standley, 2013).

### 2.1 CircRNA Detection

STARChip detects high quality circRNA by drawing power from all available samples, using multithreading to achieve rapid run times. In order to accommodate different use cases, STARChip circRNA detection can be run distributed or locally. Users provide the location of FASTQ or STAR output directories and use a parameters file to specify the minimum required reads of support and the minimum samples needed to call a circRNA. STARChip will run on paired- or single-end data of any length that STAR can align. Greater power and precision can be expected from paired-end and longer reads due to improved alignments.

#### 2.1.1 Detection and Filtration of circRNA

The initial step of in circRNA detection is searching all chimeric alignments for “back-spliced” reads. These are reads for which two chimeric segments align on the same chromosome and strand, with the 5’ segment aligning downstream of the 3’ segment (Figure 1). By default, STARChip limits circRNA to chimeric alignments 100,000 bp, chimeric junctions with less than 6 base pairs of identical sequence on each side of the junction, and does not call circRNA on mitochondrial chromosomes. CircRNA reads passing these filters are then merged if junction ends are very close (default 5bp). Each read is checked to ensure the entire alignment is within the proposed circRNA. CircRNA with less than 95% of reads aligning completely within the circRNA are eliminated as likely non-circular. Paired-end data with large inserts between reads are particularly powered to leverage this filter. CircRNA that are present in sufficient read counts and samples are carried forward for annotation and analysis. The filter requiring circRNA to be present in a minimum number of patients leverages power from all samples by removing putative circRNA detected in very few samples. An optional step is re-alignment of FASTQ files, including in the reference genome an artificial chromosome composed of circRNA sequence. This circRNA FASTA sequence is centered on the back-splice site, allowing circRNA reads to align linearly to the circRNA reference. This allows an additional strand imbalance filter. CircRNA with 10x more reads on one strand in at least 50% of samples are removed as likely false-positives.

#### 2.1.2 Quantification of circRNA

STARChip is able to quantify circRNA in two ways: the default method is to count all chimeric reads that have aligned to the filtered circRNA. An optional second method is to perform realignment of FASTQ files as described above. This strategy improves the ability to detect circRNA, especially with short and single-end reads.

#### 2.1.3 Annotation of circRNA

CircRNA are annotated for known genes, including if the junction is within an exon, intron, or outside of any known gene. STARChip automatically constructs a heat map, principle components plots, and provides a total circular read adjusted counts-per-million values.

STARChip optionally performs in-depth analysis of linear splice junctions within and around circRNA. Internal splice junctions are counted in all individuals and summarized across the entire group. This provides annotation of exons in bed format for simple assessment of the spliced size and sequence of the circRNA. Extra-circRNA linear splices are also quantified, for example forward-splices into the circRNA splice acceptor, out of the circRNA splice donor, or forward splices which envelop the circRNA by splicing from 5’ of the circRNA to 3’ of the circRNA. These measures can be useful for questions into the nature of circRNA and the genomic environments associated with circRNA formation. All linear splices in the vicinity of circRNA are summarized for each sample with a maximum linear splice value, these are useful for estimating the fraction of reads derived from circRNA at a given loci.

The output of STARChip circRNA detection is designed to be easily understandable, simple to export to additional pipelines or software, and have complex results available for interested researchers. Default settings should work in most studies, however many parameters allow the software to be completely customizable.

### 2.2 Fusion Detection

STARChip detects fusion transcripts in a distributed fashion, with each sample run separately. This allows rapid completion of large cohorts with a computing cluster. Many of the false positives common in fusion detection can be eliminated with detailed annotation of the genome. Read depth filters can be manually set by the user, or selected automatically for each sample.

#### 2.2.1 Fusion Read Support Thresholds

Automatic thresholds for required read support were developed for users to quickly and easily analyze their data. To select thresholds, we performed fusion detection with minimal read support requirements using all datasets (See Section 2.3.2). We then calculated sensitivity and the total false-positives (fusions detected in healthy tissues) as read requirements were increased. The results of this process are presented in Supplemental Figure 1. We selected as default a threshold that provides 32% sensitivity with only 15 fusions called across all healthy tissues (0.28 fusion reads per million mapped reads). The high-sensitivity threshold requires only 0.05 fusion reads per million mapped reads, which results in 42% sensitivity and 111 fusions detected in healthy tissues.

#### 2.2.2 Fusion Filtration on Read Support

Fusion detection is initialized by broadly filtering out chimeric alignments with identical sequences on each side of the junction site. Fusions mapping to the mitochondrial chromosome or an unplaced contig are filtered. Any reads with both chimeric partners mapped to known antibody parts regions are eliminated. All reads passing these filters are then joined based on location.

Putative fusions with enough read support are carried forward for further filtering. Same chromosome proximal fusions are optionally removed (these may be “read- through” transcripts that represent a different class of event than chromosomal rearrangements). Fusions are filtered for read support strand imbalance (E.g. far more reads mapping to the (+) strand than the (-) strand). If the user provides paired end data, STARChip searches for paired end reads that support the fusion without crossing the fusion junction, providing additional power to detect true and false positive fusions. Finally, STARChip filters on the number of uniquely mapping reads supporting a fusion, to reduce PCR artifacts.

#### 2.2.3 Fusion Annotation Based Filtration

Annotation based filtering of fusions is next performed, describing for each fusion partner: which genes are involved, and whether the fusion sites reside in known CNVs or repetitive elements. Fusions are filtered if the fusion partners are from the same gene, or if both partners are in a known CNV region. Fusions that appear to be circRNA are filtered. If the annotated genes of a fusion are in the same gene family, this fusion is assumed to be an alignment error and filtered. STARChip filters common false positive fusion partners. For humans, this list of known false-positives was populated using a population of over 500 non-cancerous patients, sampled and sequenced at 7 different tissues(Franzen *et al.*, 2016). Common fusion partners were examined and added to known false-positives if it was determined they were unlikely to be real.

#### 2.2.4 Fusion Output

STARChip exports fusions passing these filters, in highly detailed or streamlined format for easy examination. For each fusion, all reads contributing read support across the fusion junction are aligned and the consensus sequence is generated. This sequence can be helpful for hand-checking the validity of output fusions, or experimentally quantifying the fusion. Finally, the code for generating a circos-style plot(Krzywinski *et al.*, 2009; Ying Hu, Chunhua Yan, 2015) (Supplemental Figure 2) is output for users wishing to create circular visualizations of interchromosomal connections.

### 2.3 Performance assessment of STARChip

#### 2.3.1 Assessment STARChip circRNA

To evaluate STARChip’s ability to detect and measure circRNA, two publicly available datasets were used. The first is a high read depth RNA-Seq study of human fibroblasts both with and without RNA exonuclease digestion(Jeck *et al.*, 2013). Because exonuclease treatment selectively removes linear, but not circular RNA, this data can indicate which called circRNA are truly circular as opposed to miscalled linear RNA. Five circRNA detection tools have previously been evaluated with respect to this exonuclease data set (Hansen *et al.*, 2016), with researchers using the convention that circRNA found in the normal samples are considered bona fide if they are present at 5x higher abundance in exonuclease treated samples. We use the same convention here.

Additionally, we used the data from a study of mouse neural tissues(Rybak-Wolf *et al.*, 2015) prepared with ribosomal-depleted RNA to demonstrate common usage and the annotation features of STARChip. Details of this work can be found in the Supplemental Methods

#### 2.3.2 Assessment of STARChip fusion detection

Fusion detection with STARChip was evaluated by comparing the performance of STARChip with 3 other leading fusion detection software packages when applied to 5 different studies. SOAPFuse(Jia *et al.*, 2013) and FusionCatcher(Nicorici *et al.*, 2014) were selected given their performance in recent comparison papers(Liu *et al.*, 2016; Kumar *et al.*, 2016), while STAR-Fusion (https://github.com/STAR-Fusion) was selected because it was recently developed and is also based on the STAR aligner. To compare these methods with STARChip, we used RNA-Seq from breast cancer cell lines (“Edgren”)(Edgren *et al.*, 2011; Kangaspeska *et al.*, 2012), breast cancer cell lines validated with long-read sequencing (“Weirather”)(Weirather *et al.*, 2015), a mixture of melanoma samples and cell lines (“Berger”)(Berger *et al.*, 2010), prostate cancer samples with paired normal (“Ren”)(Ren *et al.*, 2012), and healthy tissues (“Bodymap”)(BodyMap 2.0, 2014). Only paired-end samples were examined given this is a requirement for SOAPFuse. Details of these samples can be found in Table 1.

**Table 1.**
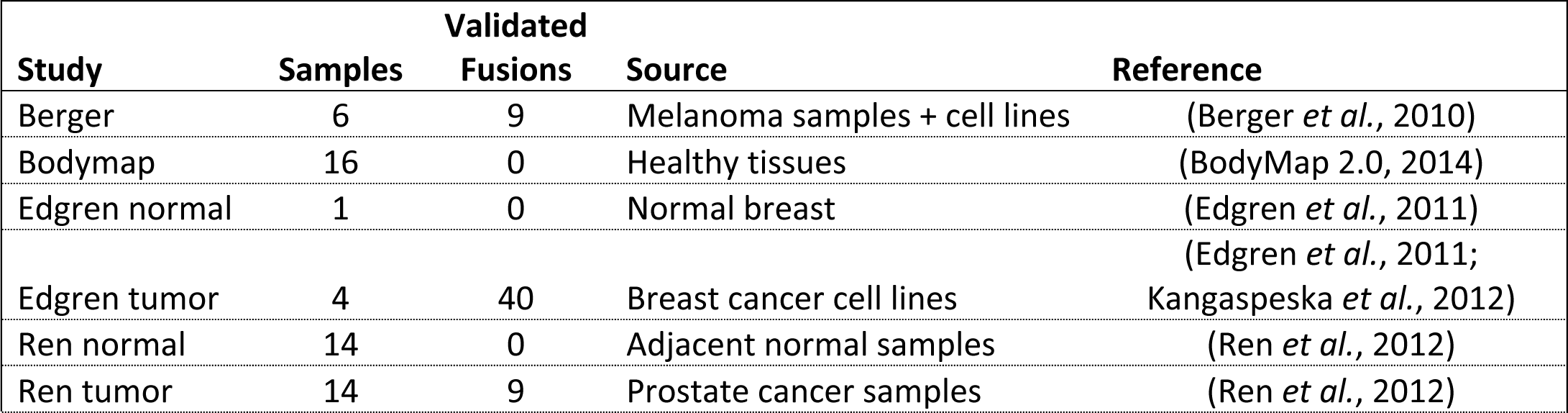
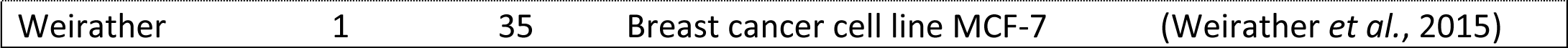
Details of Studies Used to Compare Fusion Detection Software

Samples were run using each software package on default settings. To compare time and memory usage, 64 AMD Interlagos (2.3GHz) cores on a single host with 256 GB of memory were employed. For packages supporting multithreading, parameters were set to use all 64 cores. In addition to default settings, STARChip was also run in “High Sensitivity” mode, which uses a lower reads requirement for increased sensitivity. For all software, reads were aligned to hg38, using reference gene transfer format (GTF) file Gencode v23 (http://www.gencodegenes.org) for STARChip and STAR-Fusion.

For each of Edgren, Weirather, Berger, and Ren, there are known, experimentally validated, published fusions associated with cancer tissues. Called fusions were categorized as true positives if the partner genes were identical to those of the known fusions. We calculated sensitivity as the fraction of validated fusions that could be detected. Precision was calculated as the fraction of called fusions that could be mapped to a validated fusion. It should be noted that the difficult nature of exhaustively finding and validating all fusion transcripts in cancer tissues means there exists the possibility of false-negatives in this study.

## 3. Results

We implemented STARChip to detect and quantify circRNA and fusions in several datasets. We report here the sensitivity and precision of STARChip and compare it with other leading tools in this field. Additionally we report results from STARChip analysis of new datasets.

### 3.1 STARChip circRNA

#### 3.1.1 STARChip circRNA sensitivity and precision

A previously published review of circRNA detection tools (Hansen *et al.*, 2016), assessed the effectiveness of five tools, reporting total circRNA found, and the % *bona fide* from two non-exonuclease treated samples (Jeck *et al.*, 2013). “True” circRNA were those 5x more abundant in RNAse treated samples vs untreated controls. CIRI found the most *bona fide* circRNA (2279), although this represented only 56% of all reported circRNA. Mapsplice had the best rate of *bona fide* circRNA at 73.1%, reporting 1738 true circRNA. Using these same criteria, STARChip had a 75.4% *bona fide* rate, reporting 2148 true circRNA. STARChip without circRNA index insertion and re-alignment reported a 68.7% *bona fide* rate resulting in 1124 true circRNA.

#### 3.1.2 STARChip circRNA application

The annotation and downstream application of STARChip were demonstrated using data from differently sourced mouse neural tissues, including primary cultures and cells sourced from embryonic carcinoma cell line P19 (Figure 3). Using the STARChip generated count matrix, STARChip automatically creates heatmaps (Figure 3A), and calculates principal components (Figure 3C) for quick overviews of the results. Figure 3B shows a volcano plot of circRNA comparing P19 cells to non- P19 cells, with gene IDs and splice types annotated. As a final example of STARChip facilitating downstream analysis, we merged circRNA and gene expression in a Bayesian directed network. Figure 3D shows the local regulatory network that appears to be directing the expression of ciRS-7 in these samples. This type of downstream analysis is crucial to understanding the true function of circRNA.

**Figure 3.**
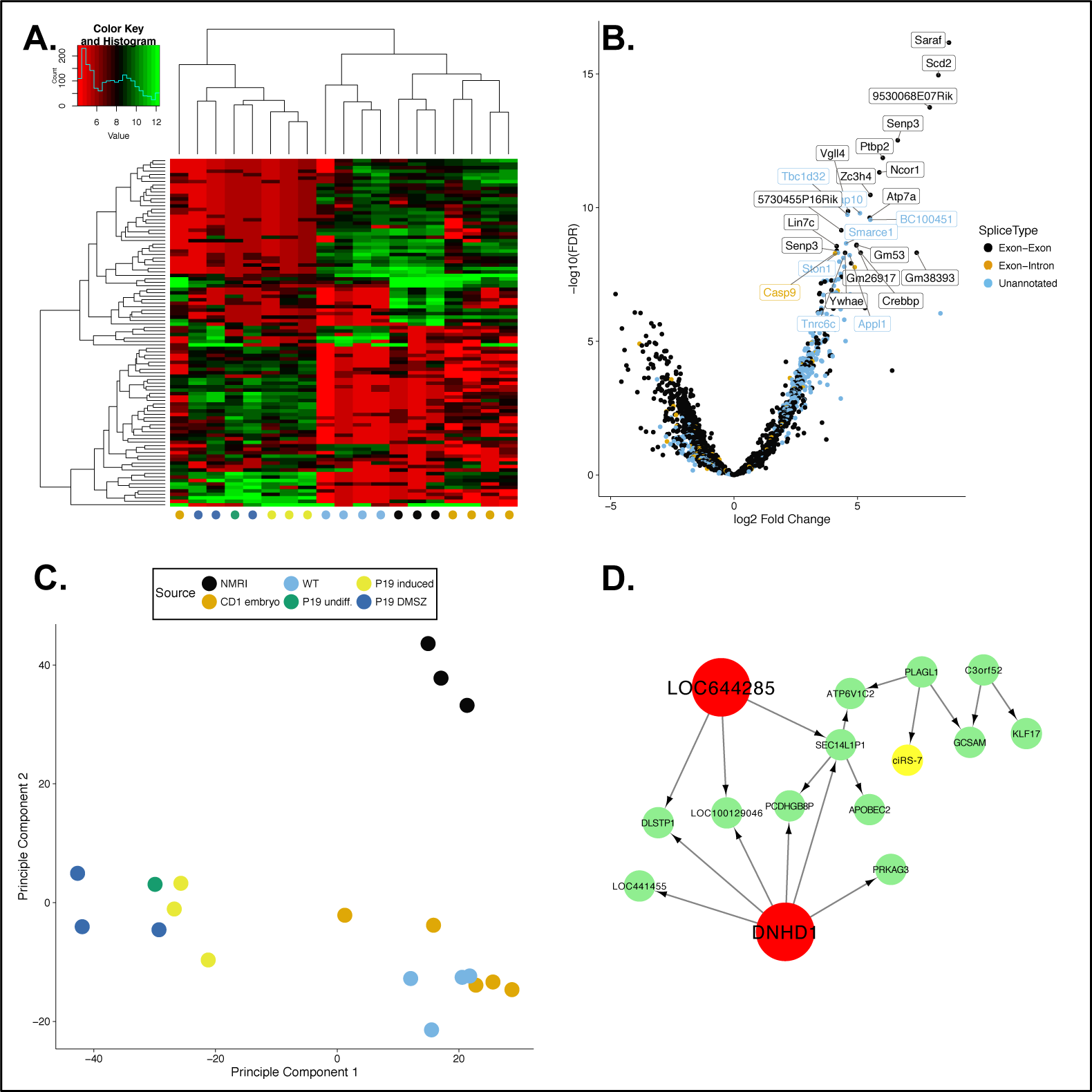
Features of STARChip circRNA analysis. A) Heatmaps are automatically generated using R, giving a visualization of the sample structure using the most variable circRNA. This figure was created using mouse samples (Rybak-Wolf *et al.*, 2015), which separate broadly along with their source. Sample groups are colored along the bottom. B) A volcano plot with each circRNA represented by a single point, where the x-axis indicates the base 2 log fold-change of P19 cells compared to non-P19 cells, and the y-axis gives the significance in false discovery rate (FDR) using a negative log base 10 scale. Individual points are labeled with the gene of origin of the circRNA and colored by the annotation classification. STARChip outputs a count matrix and circRNA annotation table, which make generating this plot simple. C) STARChip automatically carries out a principal components analysis of circRNA expression. Here three major groups are resolved, the p19 cells, wild type/CD1 induced embryonic cells, and Naval Medical Research Institute (NMRI) cells. D) STARChip output allows in-depth downstream analyses. We generated a putative directed gene coexpression network (see Supplemental Methods), which has the circRNA ciRS-7 (yellow) included. This sub-network of nearest neighbors to ciRS-7 shows genes that have been determined to be likely key drivers in red.

#### 3.1.3 STARChip circRNA runtimes

Runtimes for STARChip circRNA can vary based on the number of samples, number of computational threads used, and number of circRNA discovered. To provide some idea of computational times and memory usage, we have summarized the parameters and runtimes for the data presented here in Supplemental Table 2.

## 3.2 STARChip fusions

A guiding design principle for STARChip was improved precision, in contrast to most currently available software. This is driven by our experience with high dimensional medical omics data and the intractability of validating hundreds or even thousands of fusion calls with limited patient DNA.

### 3.2.1 STARChip fusions sensitivity and precision

Sensitivity and precision of all RNA-Seq fusion studies merged is shown in Figure 4A. STARChip is an outlier for having relatively strong precision at the expense of decreased sensitivity. Figure 4B and Table 2 show the strong variation of these values from study to study. The high precision of STARChip is again demonstrated in Figure 4C. Within two non-cancerous data sets, STARChip with default settings returned only 15 fusions, presumably false-positives. In contrast, FusionCatcher, STAR-Fusion, and SOAPFuse reported 135, 824, and 6962 fusions found in these same healthy samples.

**Figure 4.**
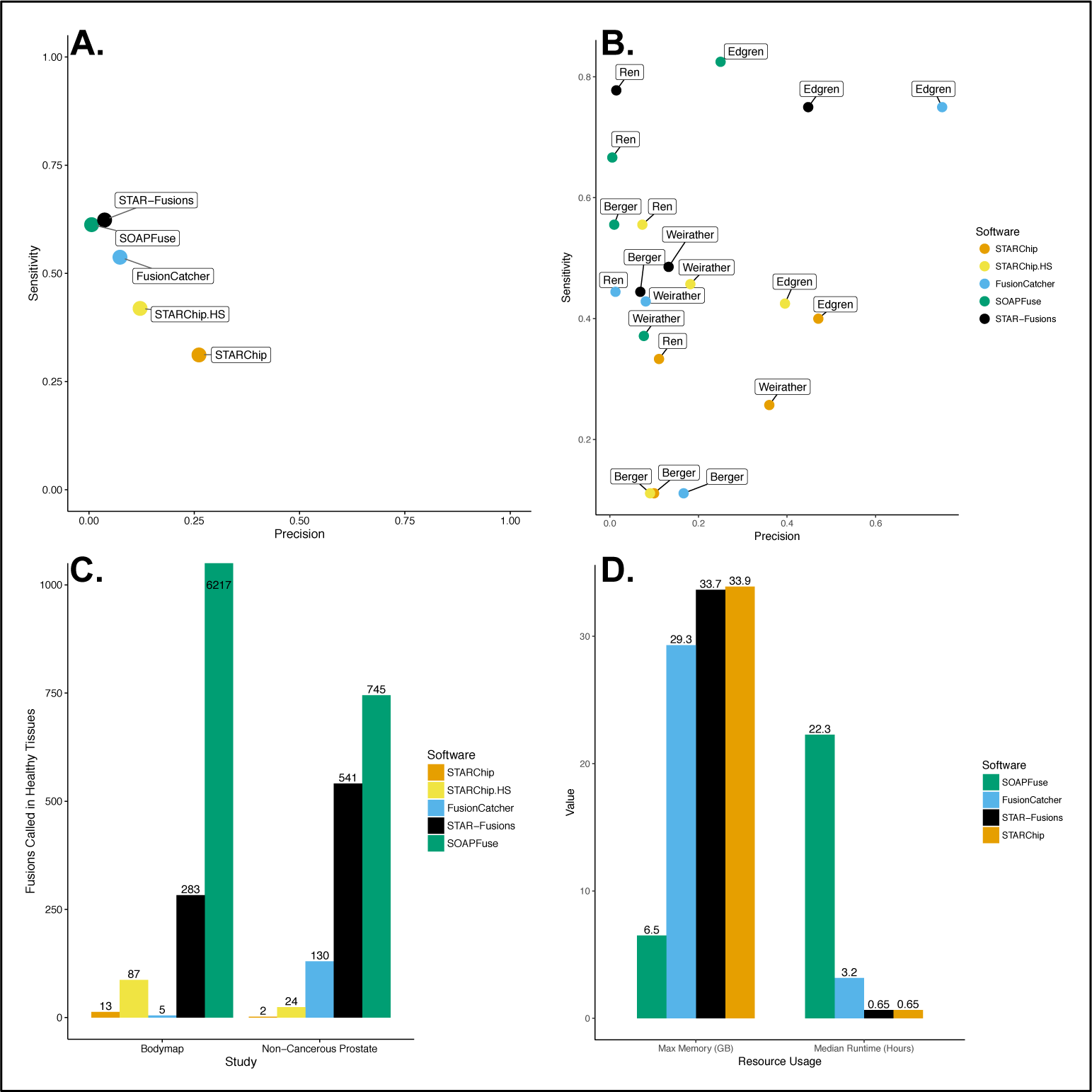
Comparison of STARChip and STARChip high sensitivity with other fusion detection software. A) Sensitivity vs. precision for all fusion-finders and all studies summarized. B) Sensitivity vs. precision separated by study. This plot demonstrates the variable efficacy of each tool by study. C) False positives are shown for two groups of healthy samples. On the left is the Illumina bodymap (BodyMap 2.0, 2014) cohort of healthy tissues, on the right are paired normal prostate samples from (Ren *et al.*, 2012). The FusionCatcher value for Bodymap is likely deflated because that software determined known false-positives using this same dataset. D) Maximum memory in gigabytes (GB) and median runtimes in hours. There was no meaningful difference in these measures for STARChip and STARChip in high sensitivity mode.

An additional benefit to STARChip is its small computational footprint. Figure 4D shows that STAR-Fusion and STARChip are extremely rapid to run, while requiring ~34 GB of memory for human or mouse genomes. The median runtimes for SOAPFuse and FusionCatcher were 22.3 hours and 3.2 hours, compared to 0.65 hours for both STAR based aligners.

## 4. Discussion

Our results indicate that in the young field of software for circRNA and fusion RNA analysis, STARChip provides key advances in fusion precision, circRNA quantification, circRNA annotation, and computational burden.

### 4.1 STARChip circRNA

With the field of circRNA research in its infancy, there are few “gold standard” datasets with which to assess the effectiveness of new tools. Individual predicted circRNA are easily validated in the laboratory, however this strategy is not feasible transcriptome-wide. Assessing confidence in discovered circRNA is often dependent on exonuclease treatment, with the expectation that circRNA will be enriched in exonuclease treated RNA, compared to non-treated samples. Although this strategy has several untested assumptions, we will proceed with it for lack of alternatives.

STARChip detects circRNA quickly by only assessing chimeric output from STAR alignments. Optionally, fastq files may be realigned with circRNA sequences inserted into the genome. The first strategy performs competitively, discovering 1124 true circRNA at a rate of 68.7% *bona fide*. Compared with other tools reviewed in (Hansen *et al.*, 2016), this is high accuracy and low sensitivity. However, with re-alignment, STARChip reports 2148 true circRNA at a rate of 75.4% *bona fide*. These values indicate better accuracy and the second highest sensitivity of the tools compared in (Hansen *et al.*, 2016), representing an improvement in overall performance of circRNA detection.

STARChip additionally attempts to streamline downstream analysis by providing high quality circRNA annotations. Gene annotations, internal splicing structure predictions, circRNA genomic size, spliced size, and alignment scores are provided for researchers to mine into the features of their circRNA of interest. This facilitates in-depth translation of circRNA counts into meaningful biological findings.

### 4.2 STARChip fusions

Ideally, RNA-Seq fusion detection would be rapid, sensitive, and precise. SOAPFuse, FusionCatcher, and STAR-Fusion all have strong sensitivity and precision values for the Edgren study. These values however, do not appear to be predictive of performance in other studies (Figure 4b). The landmark Edgren study represents the earliest available comprehensive RNA-Seq fusion data set, and most, if not all, fusion detection software is written based on the features of the fusions identified in this study. Additionally, some of the validated fusions in Edgren are present at very low read depths in the RNA-Seq data. Software tuned to detect these fusions must be hypersensitive. Unfortunately, for all other data sets, this hypersensitivity naturally results in extremely low precision values. With STARChip, we have attempted to emphasize precision at the expense of sensitivity in these particular gold-standard studies, reasoning that such hyper-tuning inflates Type I error in mining novel datasets.

The value of this strategy is emphasized in Figure 4C. For two studies of healthy tissues, the total number of fusions reported by STARChip is far lower than the other tools. An important caveat to this is that FusionCatcher used the same Bodymap data to identify false-positive fusion partners, which it automatically hard-filters. This dramatically lowers the number of fusions called in Bodymap by FusionCatcher, by design, compared to the independent data set from Ren. Fusions called from RNA-Seq must be validated in the lab using Sanger sequencing or other methods. By dramatically lowering the number of false-positive fusions called, STARChip generates output that can reasonably be tested by laboratories with modest resources.

Researchers with limited computing resources may select SOAPFuse for its ability to run on a basic workstation or laptop (6.5 GB memory), though the runtimes are much higher. With sufficient memory however, these STAR based aligners can save significant computing time. It should be noted that the majority of the time and memory requirements for STARChip arise from the STAR alignment. This alignment is run in order to facilitate expression quantitation in almost all studies that generate RNA-Seq data. Thus in the context of a typical study, running STARChip fusion detection represents a minor addition to usual computational requirements, often less than 5 minutes per sample.

Using STARChip in a pan-cancer dataset, we observed both previously reported and novel fusion events (See Supplemental Results and Discussion). These findings demonstrate the utility of STARChip in large-scale fusion screening.

## Acknowledgements

We gratefully acknowledge useful discussions with Gabriel Hoffman, Johan Bjorkegren, Lesca Holt, and Daniel Teupser.

This work was supported in part through the computational resources and staff expertise provided by Scientific Computing at the Icahn School of Medicine at Mount Sinai.

## References

Bachmayr-Heyda,A. et al. (2015) Correlation of circular RNA abundance with proliferation – exemplified with colorectal and ovarian cancer, idiopathic lung fibrosis, and normal human tissues. Sci. Rep., 5, 8057.

Berger,M.F. et al. (2010) Integrative analysis of the melanoma transcriptome. Genome Res., 1, 413–427. BodyMap 2.0 (2014).

Dobin,A. et al. (2013) STAR: ultrafast universal RNA-seq aligner. Bioinformatics, 1, 15–21.

Edgren,H. et al. (2011) Identification of fusion genes in breast cancer by paired-end RNA-sequencing. Genome Biol., 1, R6.

Franzen,O. et al. (2016) Cardiometabolic risk loci share downstream cis-and trans-gene regulation across tissues and diseases. Science, 1, 827–830.

Gao,Y. et al. (2015) CIRI: an efficient and unbiased algorithm for de novo circular RNA identification. Genome Biol., 1, 4.

Guo,J.U. et al. (2014) Expanded identification and characterization of mammalian circular RNAs. Genome Biol., 15.

Hansen,T.B. et al. (2016) Comparison of circular RNA prediction tools. Nucleic Acids Res., 1, e58–e58.

Hansen,T.B. et al. (2013) Natural RNA circles function as efficient microRNA sponges. Nature, 1, 384–388.

Jeck,W.R. et al. (2013) Circular RNAs are abundant, conserved, and associated with ALU repeats. RNA, 1, 141–157.

Jia,W. et al. (2013) SOAPfuse: an algorithm for identifying fusion transcripts from paired-end RNA-Seq data. Genome Biol., 1, R12.

Kangaspeska,S. et al. (2012) Reanalysis of RNA-Sequencing Data Reveals Several Additional Fusion Genes with Multiple Isoforms. PLoS ONE, 1, e48745.

Katoh,K. and Standley,D.M. (2013) MAFFT Multiple Sequence Alignment Software Version 7: Improvements in Performance and Usability. Mol. Biol. Evol., 1, 772–780.

Kim,D. and Salzberg,S.L. (2011) TopHat-Fusion: an algorithm for discovery of novel fusion transcripts. Genome Biol., 1, R72.

Krzywinski,M. et al. (2009) Circos: An information aesthetic for comparative genomics. Genome Res., 1, 1639–1645.

Kumar,S. et al. (2016) Comparative assessment of methods for the fusion transcripts detection from RNA-Seq data. Sci. Rep., 1, 21597.

Li,H. et al. (2009) The Sequence Alignment/Map format and SAMtools. Bioinforma. Oxf. Engl., 1, 2078–2079.

Liu,S. et al. (2016) Comprehensive evaluation of fusion transcript detection algorithms and a meta-caller to combine top performing methods in paired-end RNA-seq data. Nucleic Acids Res., 1, e47–e47.

Mertens,F. et al. (2015) The emerging complexity of gene fusions in cancer. Nat. Rev.Cancer, 1, 371–381.

Nicorici,D. et al. (2014) FusionCatcher – a tool for finding somatic fusion genes in paired-end RNA-sequencing data.

Quinlan,A.R. (2014) BEDTools: The Swiss-Army Tool for Genome Feature Analysis: BEDTools: the Swiss-Army Tool for Genome Feature Analysis. In, Bateman,A. et al. (eds), Current Protocols in Bioinformatics. John Wiley & Sons, Inc., Hoboken, NJ, USA, p. 11.12.1–11.12.34.

R Core Team (2015) R: A language and environment for statistical computing. R Foundation for Statistical Computing, Vienna, Austria.

Ren,S. et al. (2012) RNA-seq analysis of prostate cancer in the Chinese population identifies recurrent gene fusions, cancer-associated long noncoding RNAs and aberrant alternative splicings. Cell Res., 1, 806–821.

Rybak-Wolf,A. et al. (2015) Circular RNAs in the Mammalian Brain Are Highly Abundant, Conserved, and Dynamically Expressed. Mol. Cell, 1, 870–885.

Salzman,J. et al. (2013) Cell-Type Specific Features of Circular RNA Expression. PLoS Genet., 1, e1003777. STAR-Fusion/STAR-Fusion GitHub.

Wang,P.L. et al. (2014) Circular RNA Is Expressed across the Eukaryotic Tree of Life. PLoS ONE, 1, e90859.

Weirather,J.L. et al. (2015) Characterization of fusion genes and the significantly expressed fusion isoforms in breast cancer by hybrid sequencing. Nucleic Acids Res, 1, e116–e116.

Ying Hu, Chunhua Yan (2015) OmicCircos: High-quality circular visualization of omics data.

